# Identification of a Sulf2-dependant astrocyte subtype that stands out through the expression of Olig2 in the ventral spinal cord

**DOI:** 10.1101/430074

**Authors:** David Ohayon, Nathalie Escalas, Philippe Cochard, Bruno Glise, Cathy Danesin, Cathy Soula

## Abstract

During spinal cord development, both spatial and temporal mechanisms operate to generate glial cell diversity. Here, we addressed the role of the Heparan Sulfate-editing enzyme Sulf2 in the control of gliogenesis in the mouse developing spinal cord and found an unanticipated function for this enzyme. Sulf2 is expressed in ventral spinal progenitors at initiation of gliogenesis, including in Olig2-expressing cells of the pMN domain known to generate most spinal cord oligodendrocyte precursor cells (OPCs). We found that Sulf2 is dispensable for OPC development but required for proper generation of an as-yet-unidentified astrocyte precursor cell (AP) subtype. These cells, like OPCs, express Olig2 while populating the spinal parenchyma at embryonic stages but also retain Olig2 expression as they differentiate into mature astrocytes. We therefore identify a spinal Olig2-expressing AP subtype that segregates early under the influence of the extracellular enzyme Sulf2.

## Introduction

Glial cells are recognized for serving critical functions in the central nervous system (CNS). Among them, astrocytes and oligodendrocytes, the neural derivatives, populate all regions of the mature CNS. Astrocytes are active regulators of synaptogenesis and neurotransmission and are also critical for formation of the blood-brain barrier. The mostly known feature of oligodendrocytes is formation of myelin sheaths that provides insulation of neuronal axons.

Acquisition of these two distinct glial cell fates is acknowledged to result from spatial segregation of neural progenitors that is established early, during patterning of the neural tube. A good understanding of organizing signals and target genes involved in this process was acquired studying the development of the embryonic spinal cord (Dessaud et al., 2008). Under the influence of patterning signals, progenitors of the ventral neural tube activate or repress sets of transcription factors that subsequently subdivide it into distinct progenitor domains, each dedicated to first generate specific neuronal subtypes, followed by glial cells (Ben Haim and Rowitch, 2017). Oligodendrocyte precursor cells (OPCs) originate from the pMN domain that earlier on produce motor neurons (MNs) while three ventral astrocyte precursor cell (AP) subtypes (VA1–3) are generated from p1-p3 domains that previously generate V1–3 interneurons. Thus, the origins of OPCs and APs are considered to be more closely connected to neuron subtype progenitors than with each other. The mechanistic parallel between neuron and glial cell development is well exemplified by functional analysis of the transcription factor Olig2, expressed in pMN cells. In Olig2 null spinal cords, pMN cells fail to generate both MNs and OPCs (Lu et al., 2002; Takebayashi et al., 2002; Zhou and Anderson, 2002). Instead, in these mutants, supernumerary V2 interneurons and APs are produced as a consequence of transformation of the presumptive pMN domain into a p2 domain. Therefore, spinal OPCs and APs have been proposed to develop along mutually exclusive paths, Olig2 being viewed as a repressor of a ‘pro-astrocytic’ program (Rowitch and Kriegstein, 2010). Olig2 is also considered as a key determinant of OPC specification, maturation and differentiation and it continues to be present in differentiated oligodendrocytes of the adult spinal cord (Meijer et al., 2012). However, contrasting with the notion that Olig2 represses the astrocytic fate, lineage-tracing studies brought evidence that Olig2-expressing progenitors of the pMN domain generate some astrocytes (Masahira et al., 2006; Tsai et al., 2012). Furthermore, Olig2 expression has been reported in grey matter astrocytes of the adult mouse spinal cord (Barnabé-Heider et al., 2010; Guo et al., 2011).

Beyond spatial control, a temporal component is important to initiate gliogenesis. Although timing mechanism regulating AP commitment remains elusive, OPC commitment from pMN cells is known to rely on a temporal rise of Sonic Hedgehog (Shh) signalling activity (Danesin and Soula, 2017). The heparan sulfate–editing enzyme Sulf1, known to modulate the sulfation state of heparan sulfate proteoglycans (HSPGs) at the cell surface (Lamanna et al., 2007), is a key player in this process. This enzyme is expressed specifically by Shh-secreting cells of the floor plate immediately prior to OPC specification. Sulf1, by eliminating 6O sulfate groups on HS chains, locally lowers Shh/HSPG interaction and thereby promotes Shh release at the right time to trigger OPC specification from Olig2-expressing pMN cells (Touahri et al., 2012; Al Oustah et al., 2014). Jiang and collaborators (2017) more recently reported that Sulf2, the second member of the Sulf protein family (El Masri et al., 2017), plays a similar role as Sulf1 in triggering the MN to OPC fate change. In the present work, we also investigated the function of Sulf2 in the control of gliogenesis and came to the different conclusion that Sulf2 is dispensable for OPC production. Instead, we found that Sulf2, which in contrast to Sulf1 is expressed in Olig2-expressing pMN cells, promotes generation of a discrete and previously uncharacterized subtype of ventral spinal cord APs distinguishable from others by the expression of Olig2.

## Results

### Sulf2 controls generation of glial precursors marked by Olig2 expression but distinct from OPCs

In the mouse developing spinal cord, Olig2-expressing (Olig2+) pMN cells start generating OPCs from 12.5 days of development (E12.5). At this initial stage of gliogenesis, *sulf2* is expressed in ventral progenitors, including in Olig2+ cells of the pMN domain (Figure 1A). Noticeably, distinct levels of *sulf2* mRNA staining were observed in this domain (Figure 1a-a’), indicating heterogeneous levels of *sulf2* expression within the population of Olig2+ pMN cells. Later on, *sulf2* expression is maintained in ventral progenitors but also in few cells streaming away in the spinal parenchyma (Figure 1B). At E14.5, a subset of *sulf2+* parenchymal cells retain expression of Olig2 (Figure 1B-b’). Based on these data, Sulf2 appeared to be a reasonable candidate to play a role in OPC development. To test this possibility, we analyzed OPC generation in *sulf2* knocked-out (*sulf2-/-*) embryos using Olig2 together with Sox10, a reliable marker of oligodendroglial cells which is up-regulated in Olig2+ pMN cells as soon as they commit to the OPC fate (Zhou et al., 2000). As previously reported (Touahri et al., 2012), we found that, at E12.5 in wild-type embryos, almost 50% of Olig2+ pMN cells coexpress Sox10 (Figure 1C, E, F). Few Olig2+ cells that have already emigrated in the spinal parenchyma were also observed and all of them were found coexpressing Sox10 (Figure 1C). Similar pattern of Olig2 and Sox10 expression was observed in *sulf2-/-* littermates (Figure 1D) and cell counting indicated that the same number of Olig2+/Sox10+ OPCs has been generated at E12.5 in *sulf2-/-* and wild-type embryos (Figure 1E, F). Therefore, contrasting with previous study (Jiang et al., 2017), our data indicated that Sulf2 is dispensable for initiation of OPC production in the ventral spinal cord. To get further insight on this disparity of results, we next considered the possibility that Sulf2 might impair ongoing OPC production at later developmental stages. We then examined Olig2 and Sox10 expression one day later, at E13.5, and, again, found similar pattern of Olig2 and Sox10 expression in *sulf2-/-* and wild-type littermate embryos (Figure 1G-H’’, K). We therefore concluded that Sulf2 activity is not required to sustain OPC generation. However, we made at that stage a quite unexpected observation. We found that, in wild-type embryos, while coexpression of Olig2 and Sox10 was detected in most cells streaming away from the pMN domain, expression of Sox10 remained undetectable in a subset of parenchymal Olig2+ cells (Figure 1G-G’’, I-I’’). These Olig2+/Sox10^-^ cells were predominantly found close to the progenitor zone, suggesting that they have been recently generated. In support of this, we never detected Olig2+/Sox10^-^ cells in the parenchyma at E12.5 (Figure 1C). Strikingly, only few Olig2+/Sox10^-^ parenchymal cells were detected in *sulf2-/-* littermates (Figure 1H-H’’, JJ’’) and quantification of these cells showed that their number was significantly reduced in *sulf2-/-* embryos compared to wild-type littermates (Figure 1L). Deficient generation of Olig2+/Sox10^-^ cells but not of Olig2+/Sox10+ OPCs was still apparent at fetal stage E18.5, where Olig2+/Sox10^-^ cells were still found populating the ventral spinal cord (Figure S1A-D). These results, showing that the number of Olig2+/Sox10^-^ cells but not of Olig2+/Sox10+ cells was reduced in *sulf2-/-* embryos, clearly pointed out to a specific function of *sulf2* in controlling generation of only a subtype of Olig2+ parenchymal cells.

**Figure 1:**
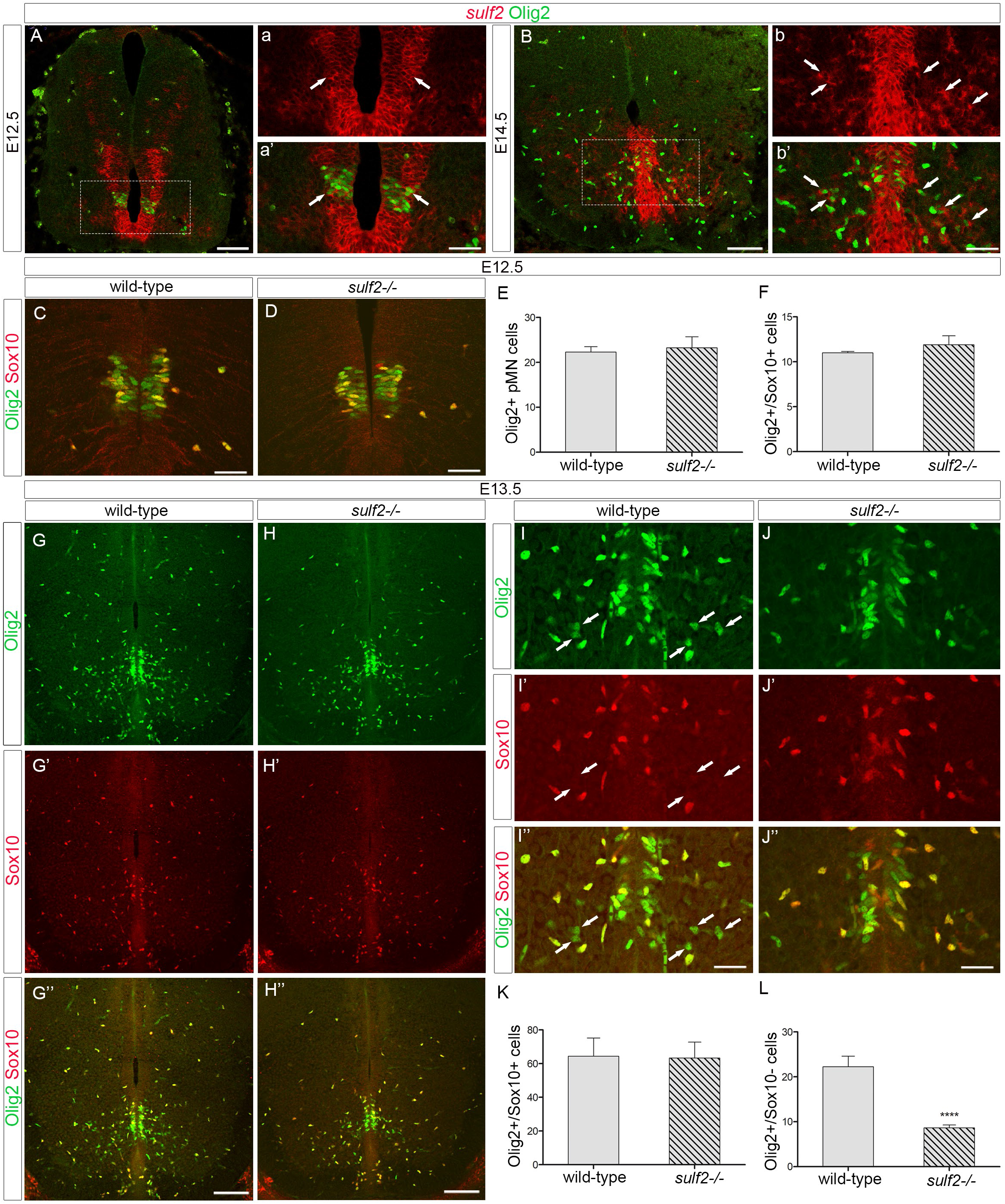
Sulf2 depletion impairs production of Olig2-expressing precursors distinct from OPCs. All images show transverse spinal cord sections. **A-b’**: Combined detection of *sulf2* mRNA (red) and Olig2 (green) at E12.5 (A-a’) and E14.5 (B-b’). a,a’ and b,b’ show higher magnification of the area framed in A and B, respectively. **C, D, G-J’’**: Double detection of Olig2 (green) and Sox10 (red) at E12.5 (C, D) and E13.5 (G-J’’) in wild-type **(C, G-G’, I-I’’)** and *sulf2-/-* (D, H-H’’, J-J’’) embryos. Note reduced number of Olig2+/Sox10^-^ cells in the spinal parenchyma of *sulf2-/-* (H’’, J’’) compared to wild-type (G’’, J”) E13.5 embryos. **E-F**: Quantification of Olig2+ pMN cells (E) and Olig2+/Sox10+ cells (F) in E12.5 wild-type and *sulf2-/-* embryos. **K, L**: Quantification of Olig2+/Sox10+ cells (K) and Olig2+/Sox10^-^ cells (L) in the E13.5 spinal parenchyma of wild-type and *sulf2-/-* embryos. Data are presented as mean ± SEM (**** p ˂ 0.0001). Scale bars = 100 μm in A, B, G-H’’ and 50 μm in a, a’, b, b’, C, D, I-J’’. See also Figure S1.

We next asked whether Sulf2 expression specifically in Olig2+ cells is required for proper generation of the Olig2+/Sox10^-^ cell subtype. We deleted *sulf2* in Olig2+ cells by breeding *sulf2* floxed mice (*sulf2fl/fl*, Tran et al., 2012) with *olig2* promoter-driven Cre (*Olig2-cre*) knock-in mice (Dessaud et al., 2007). Counting of Olig2+/Sox10+ (OPCs) and Olig2+/Sox10^-^ cells was performed in (*Olig2-cre*^+/−^*;sulf2fl/fl*) embryos and in control wild-type, *Olig2*-*Cre*^+/−^ and (*Olig2*-*Cre*^+/−^*/sulf2fl/+*) littermates at E14.5 and E16.5 (Figure S1E-P). As previously reported (Liu et al., 2007), we found that loss of one copy of *olig2*, as is the case in *Olig2-Cre*^+/−^ embryos, impairs generation of Olig2+/Sox10+ OPCs compared to wild-type littermates. Similarly, Olig2+/Sox10^-^ cells were reduced in number in *Olig2*-*Cre*^+/−^ compared to wild-type embryos, indicating a dosage-dependent regulation by Olig2 also for Olig2+/Sox10^-^ development. We found no significant difference in cell numbers between *Olig2*-*Cre*^+/−^ and *Olig2*-*Cre*^+/−^*/sulf2fl/+* embryos. By contrast, the number Olig2+/Sox10^-^ cells appeared significantly reduced in *Olig2-cre*^+/−^*;sulf2fl/fl* embryos compared to *Olig2-Cre*^+/−^*/sulf2fl/+* littermates while no difference was found in the numbers of Olig2+/Sox10+ OPCs. Therefore, Sulf2 depletion specifically in Olig2+ cells is sufficient to impair generation of Olig2+/Sox10^-^ cells.

Together, our data point out two important issues: i) two molecularly distinct Olig2+ precursor subtypes, namely the Olig2+/Sox10+ OPCs and a second cell population expressing Olig2 but not Sox10, are generated in the ventral spinal cord, ii) Sulf2 expression by Olig2+ cells specifically controls generation of the Olig2+/Sox10^-^ cell subtype. For convenience, we refer to this sub-population henceforth as Sulf2-dependant Olig2+ cells.

### Sulf2-dependant Olig2+ cells express astroglial identity genes

Supporting the view that Sulf2-dependant Olig2+ cells are distinct from OPCs, parenchymal Olig2+/Sox10^-^ cells neither expressed PDGFRα nor Olig1 (Figure 2A-B), two specific hallmarks of OPCs in the spinal cord (Pringle and Richardson, 1993; Meijer et al., 2012). We then investigated the possibility that Olig2+/Sox10^-^ cells might belong to the astroglial lineage by combining immunodetection of Olig2 and in situ localization of *fgfr3, aldh1L1* and *tenascin-C* mRNAs, all reported to mark spinal APs as well as progenitors from which they originate (Pringle et al., 2003; Cahoy et al., 2008; Karus et al., 2011). At E13.5, *fgfr3* mRNA was detected in the progenitor zone where its expression domain overlaps the Olig2+ pMN domain (Figure 2C). At this stage, few individual *fgfr3*+ cells were also detected in the spinal parenchyma and, noticeably, most, if not all, of them were positive for the Olig2 staining (Figure 2C), indicating that the first *fgfr3*+ cells to leave the progenitor zone express Olig2. By E15.5, more *fgfr3+*/Olig2+ parenchymal cells were found but *fgfr3*+ cells that do not express Olig2 were also observed emigrating from the ventrally and dorsally located progenitor domains (Figure 2D). Similar results were obtained detecting *aldh1L1* or *tenascin-C* mRNAs and the Olig2 protein (Figure 2E, F). These data therefore revealed the existence of a ventral spinal cord AP subtype that distinguishes itself from others by Olig2 expression.

**Figure 2:**
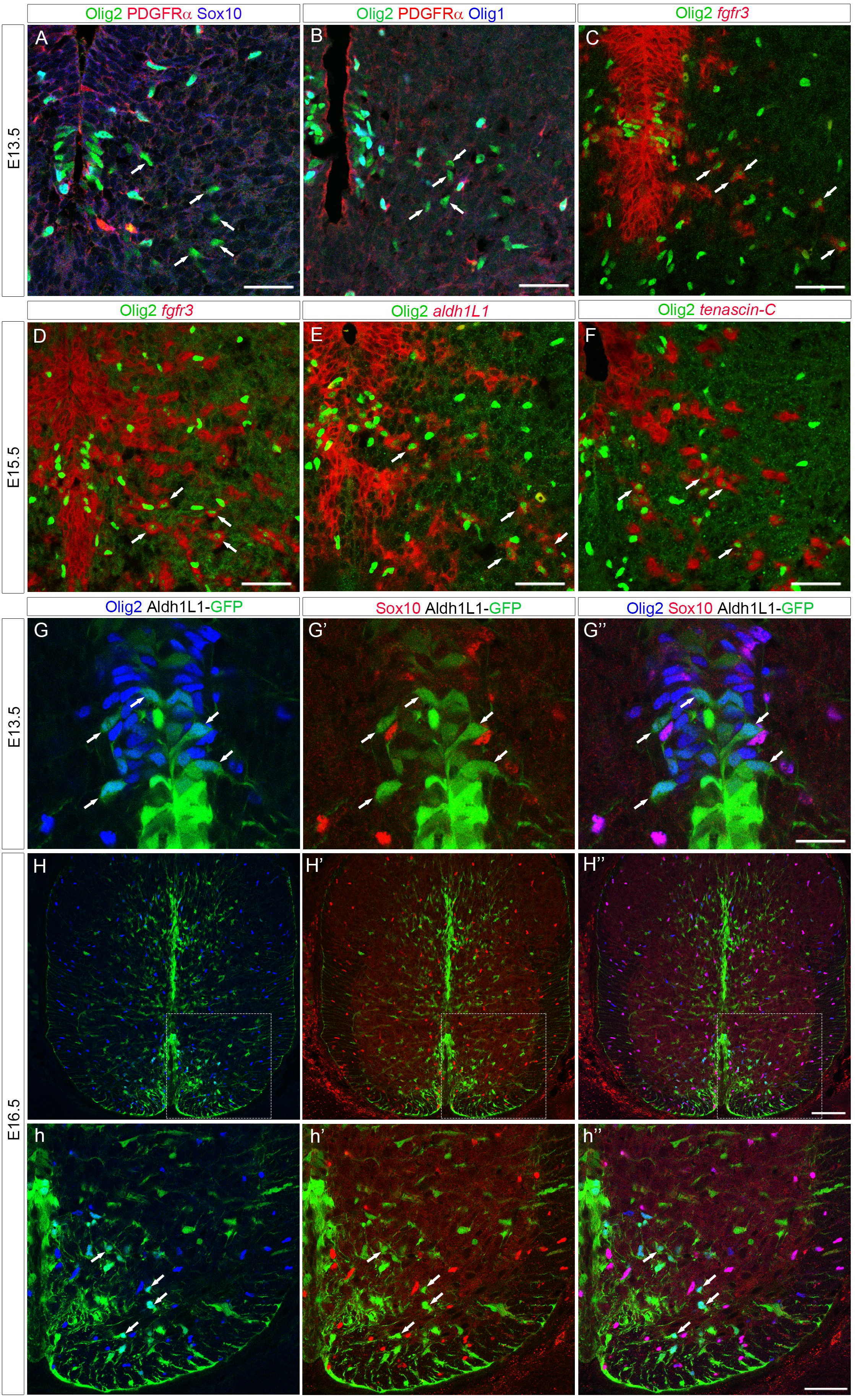
Sulf2-dependant Olig2+ cells express AP but not OPC markers. All images show transverse spinal cord sections. **A, B**: Combined detection of Olig2 (green), PDGFRα (red) and Sox10 (A, blue) or Olig1 (B, blue) viewed on E13.5 hemi-ventral spinal cord. Note the presence of Olig2+ cells that are not stained with the OPC markers (arrows). **C-F**: Combined detection of Olig2 (green) and mRNAs (red) encoding for *fgfr3* (C, D), *aldh1L1* (E) and *tenascin-C* (F) viewed on E13.5 (C) and E15.5 (D-F) hemi-ventral spinal cords. Note, in all panels, the presence of Olig2+ cells stained with mRNA probes (arrows). **G-G’’**: Combined detection of Olig2 (blue), Sox10 (red) and GFP (green) on E13.5 Aldh1L1-GFP transgenic embryo. Images show high magnification of the ventral progenitor zone. Horizontal sets present successively Olig2 staining, Sox10 staining, and the merged image. Note the presence of GFP+/Olig2+/Sox10^-^ cells (arrows) both in the progenitor zone (Olig2+ pMN domain) and in the adjacent spinal parenchyma. **H-h’’**: Combined detection of Olig2 (blue), Sox10 (red) and GFP (green) on E16.5 Aldh1L1-GFP transgenic embryo. h, h’ and h’’ show higher magnifications of the area framed in H, H’ and H’’, respectively. Arrows point to GFP+/Olig2+/Sox10^-^ cells. Scale bars = 100 μm in H-H’’, 50 μm in A-F, h-h’’ and 25 μm in G-G’’. See also Figures S2.

Due to the setback of combining Sox10 immunostaining with in situ hybridization, we were not able to directly define whether *fgfr3*+/Olig2+ APs match the Olig2+/Sox10^-^ cell population. We therefore turned to Aldh1L1-GFP astrocyte-specific reporter mice (Cahoy et al., 2008) and performed immunodetection of the reporter protein together with that of Olig2 and Sox10. We first focused on the progenitor zone at E13.5 and found the presence of Aldh1L1-GFP+/Olig2+ included in the pMN domain (Figure 2G). We also observed at this stage some early born Aldh1L1-GFP+/Olig2+ cells located in the adjacent parenchyma (Figure 2G). Comparison with the Sox10 staining allowed detecting Sox10+ OPCs both in the pMN domain and in the ventral parenchyma but Sox10 was never detected in Aldh1L1-GFP+/Olig2+ cells either located within the pMN domain or that have already emigrated from the progenitor zone (Figure 2G’,G’’). These data, by revealing a subtype of Olig2+ pMN cells expressing Aldh1L1 but not Sox10 at E13.5, support the view that the Olig2+ APs and the Sulf2-dependent Olig2+ cells represent one and the same cell population. The two Olig2+ cell subtypes, expressing either Aldh1L1-GFP or Sox10, were still detected in the spinal parenchyma at E16.5 (Figure 2 H-h’’). At this stage, Aldh1L1-GFP+/Olig2+ APs remained located within the ventral spinal parenchyma while Olig2+/Sox10+ OPCs dispersed in all directions (Figure 2 H-h’’). To learn more about the Olig2+ AP subtype, we performed immunostaining using antibodies specific for Nkx6.1, expressed in AP originating from ventral progenitors, and for NFIA and Zeb1, two transcription factors broadly expressed in newly generated spinal APs (Deneen et al., 2006; Zhao et al., 2014; Ohayon et al., 2016). We found that essentially all Olig2+/Sox10^-^ cells coexpress these proteins at E14.5 (Figure S2-C’’), indicating that they share the same basic properties as previously identified ventral APs.

Together, these data highlight a subtype of APs that emerge from Olig2-expressing progenitors and maintain Olig2 expression as they populate the ventral spinal parenchyma.

### Sulf2-dependent APs maintain expression of Olig2 while activating astrocyte differentiation genes

We next examined whether, as reported for cortical white matter APs (Marshall et al., 2005; Cai et al., 2007), expression of Olig2 in ventral spinal cord APs reflects an immature stage of astrocytogenesis. Specific markers for different stages of AP maturation are not available. However, acquisition of mature astrocyte identity in the spinal cord can be followed according to the previously reported stepwise process of their maturation (Tien et al., 2012). This process progress through different stages: a proliferating intermediate AP stage, during which parenchymal APs remain proliferative (E14.5 to P3), a maturing postnatal astrocyte stage (from P5) in which APs no more proliferate and an adult astrocyte status. We therefore assessed Olig2 and Sox10 expression at early (P2) and late (P7) post-natal stages as well as in adult spinal cord, postulating that, if Olig2 expression in Sulf2-dependent Olig2+ APs is down-regulated as these cells undergo terminal differentiation, the population of Olig2+/Sox10^-^ cells should progressively disappear. At P2, Olig2+/Sox10^-^ cells were still detected in the ventral spinal cord (Figure 3A, a’’). EdU injection over the period of P0 to P2 indicated that 20% of these cells incorporated EdU (Figure S3A-B), corresponding to the expected proliferation profile for APs at early post-natal stages (Tien et al., 2012). Thereafter and until adulthood, we detected equivalent number of Olig2+/Sox10^-^ cells in the ventral spinal cord and no significant difference was found in their rates relative to the Olig2+/Sox10+ oligodendroglial cell population (Figure 3B-D). Moreover, at post-natal and adult stages, Olig2 still showed colocalisation with the AP marker *fgfr3* (Figure S3C-F). Therefore, the Olig2+/Sox10^-^ ventral spinal cell population persists over the period of astrocyte maturation.

**Figure 3:**
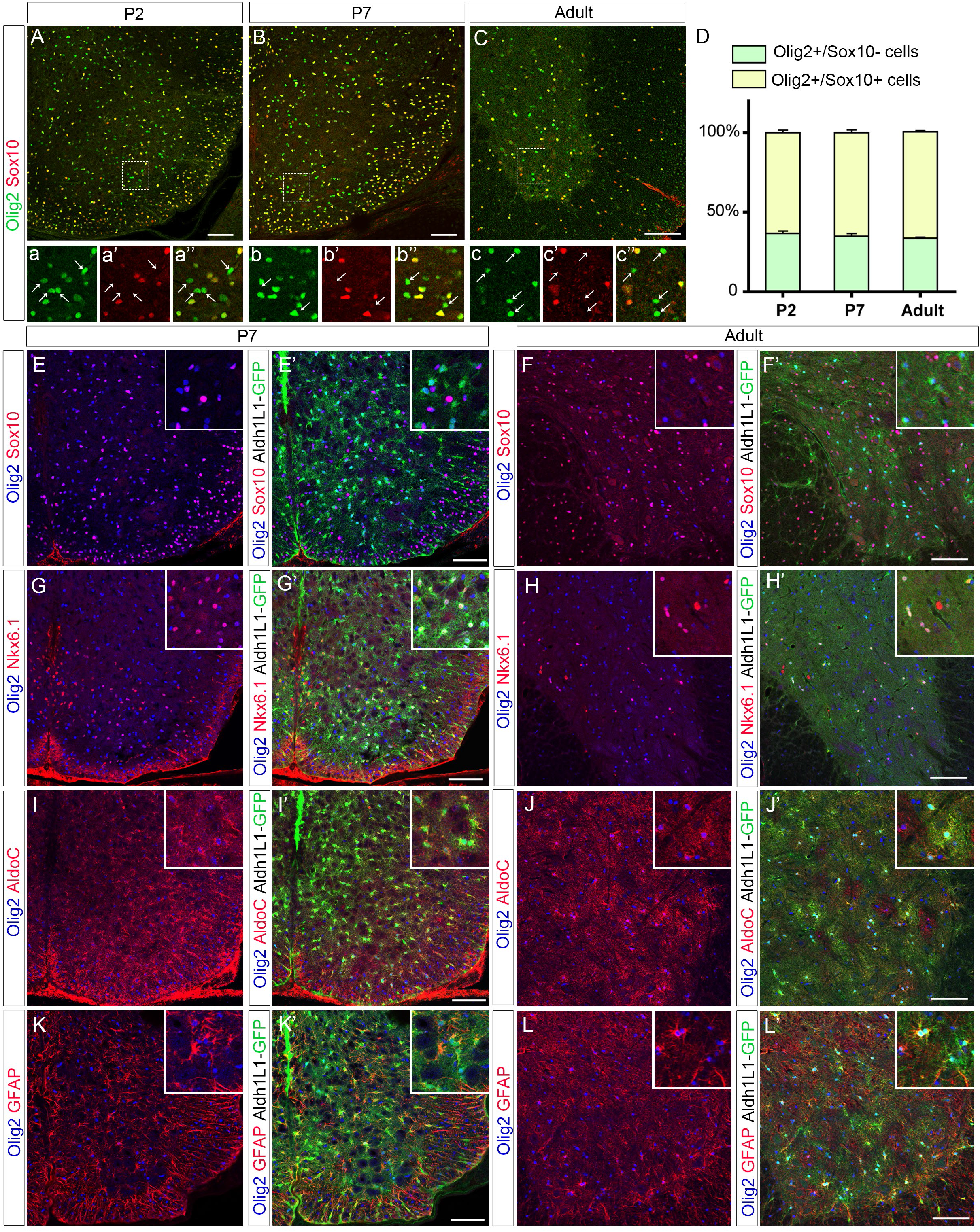
A sub-set of ventral spinal cord astrocytes retain expression of Olig2 at post-natal and adult stages. Images show transverse sections of hemi-ventral spinal cord. **A-c’’**: Double detection of Olig2 (green) and Sox10 (red) at P2 (A, a-a’’), P7 (B, b-b’’) and adult (C, c-c’’) stages. Images aa’’, b-b’’ and c-c’’ show higher magnification of the areas framed in A, B and C, respectively. Horizontal sets present successively Olig2 staining, Sox10 staining, and the merged image. Note the presence of Olig2+/Sox10^-^ cells (arrows) at all stages. **D**: Proportion of Sox10^-^ and Sox10+ cells on the Olig2+ cell population in the ventral spinal cord at P2, P7 and at adult stage. **E-L’**: Combined detection of Olig2 (blue) and Sox10 (E, F, red), Nkx6.1 (G-H, red), AldoC (I-J, red) or GFAP (K-L, red) in Aldh1L1-GFP transgenic individuals at P7 (E, G, I, K) and at adult stage (F, H, J, L). Prime panels add GFP (green) to the corresponding double immunostaining. Insets show higher magnification of the area framed in each panel. Data are presented as mean ± SEM. Scale bars = 100 μm. See also Figures S3 and S4.

We next performed characterization of Olig2+ cells in P7 and adult Aldh1L1-GFP spinal cords. We found extensive coexpression of Aldh1L1-GFP and Olig2 but not Sox10 at P7 and at adult stage (Figure 3E-F’). Supporting the view that they originate from ventral progenitors, Aldh1L1-GFP+/Olig2+ cells were all found coexpressing Nkx6.1 (Figure 3G-H’). We next examined expression of the astrocytic marker AldoC and found that a fraction of ventral Aldh1L1-GFP+/AldoC+ cells indeed express Olig2 at both stages (Figure 3I-J’). We then analyzed expression of the astrocyte maturation marker GFAP and also found Aldh1L1-GFP+/Olig2+ cells expressing GFAP in P7 and adult individuals (Figure 3K-L’). To confirm this, we analyzed expression Olig2 and Sox10 in GFAP-GFP transgenic mice in which expression of the reporter allows complete staining of astrocyte cell bodies (Nolte et al., 2001). In these mice, GFAP-GFP+ cells coexpressing Olig2 was also detected in the ventral spinal parenchyma at P7 (Figure S4D-D”).

Together, these data indicate that Olig2 expression is retained in a subtype of astrocytes as they maturate in the ventral spinal cord.

### Sulf2 depletion does not cause major spinal cord AP deficit

Sulf2 expression is not restricted to the pMN domain but extended ventrally and dorsally in progenitor domains known to generate V1 to V3 APs (Figure 1A, B). Moreover, *sulf2*+ cells were detected streaming away at all levels of the ventral progenitor zone (Figure 3A-C), within a pattern very suggestive of *sulf2* expression being maintained in ventral spinal cord APs. Supporting this, most, if not all, parenchymal *sulf2*+ cells coexpress *fgfr3* but not *sox10* at E13.5 (Figure 4D-E). We next logically asked whether Sulf2 might have a general influence on AP generation. We thus compared the time course of *fgfr3* expression in wild-type and *sulf2-/-* embryos. At E13.5, when the first wave of AP production became discernible in wild-type ventral spinal cord (Figure 4F), only very few *fgfr3*+ cells were found emigrating from the progenitor zone in *sulf2-/-* embryos (Figure 4G) and cell quantification confirmed defective production of *fgfr3*+ cells in *sulf2-/-* embryos compared to wild-type littermates (Figure 4H). Thus, as expected since the first APs to be generated are those expressing Olig2 (see Figure 2C), Sulf2 depletion causes defective AP production at initiation of astrogenesis. From E14.5, although cell counting still indicated a slight decrease in the number of ventral *fgfr3*+ cells in mutant embryos, numerous *fgfr3*+ individual cells were found populating the spinal parenchyma both in wild-type and *sulf2-/-* embryos (Figure 4I-K). At E18.5, *fgfr3*+ cells populate all regions of the spinal cord and cell counting indicated a downward trend, although no significant, of their number in *sulf2-/-* compared to wild-type individuals (Figure 4O-Q), indicating no gross effect of Sulf2 depletion on astrocyte development at this stage. However, combination of GFAP and Olig2 detection at this stage showed a significant reduction in the number of GFAP+/Olig2+ cells in *sulf2-/-* compared to wild-type embryos, confirming the requirement of Sulf2 activity for proper generation of this astrocyte subtype (Figure S4B-D).

**Figure 4:**
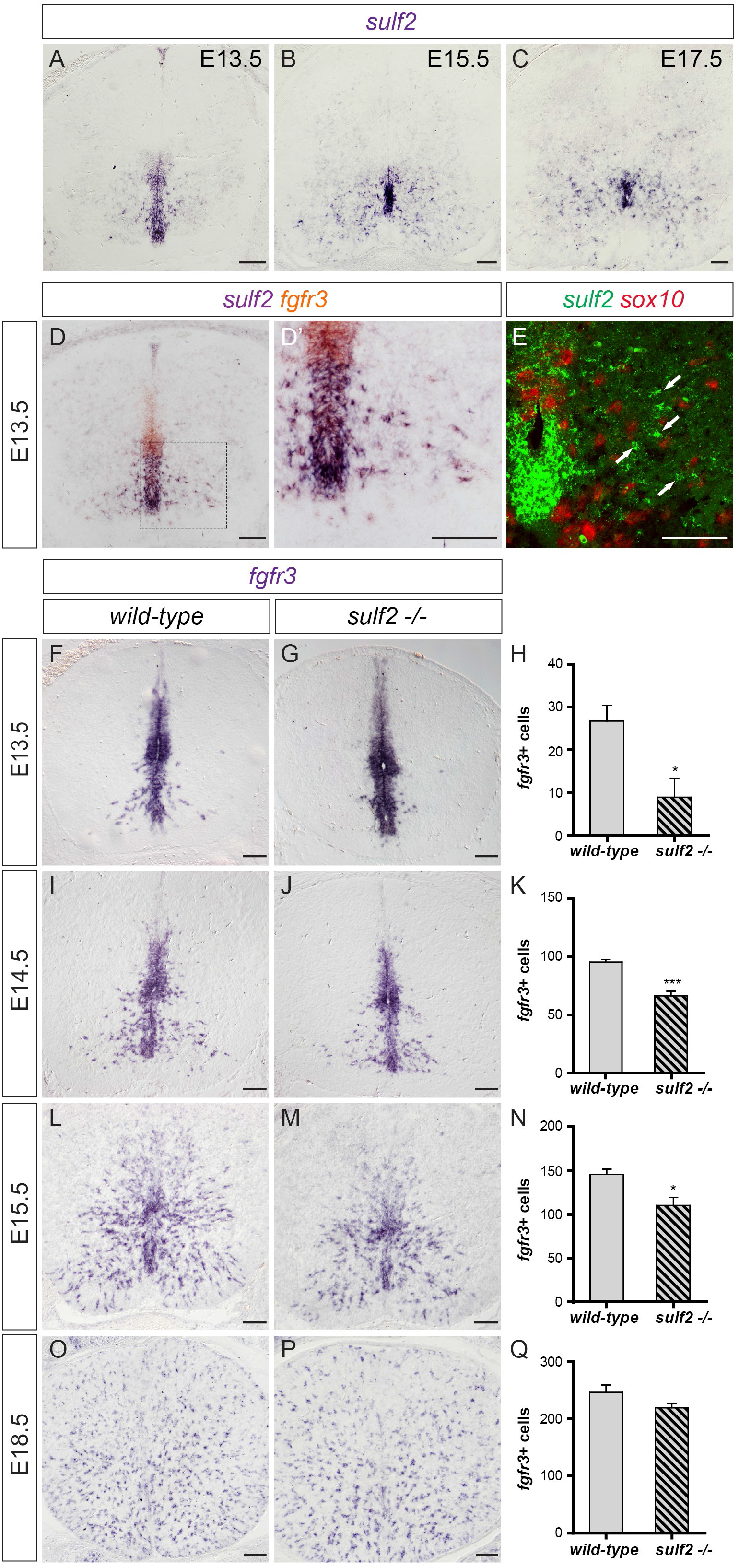
Sulf2 is widely expressed in ventral spinal APs but its depletion does not cause major defect of astrocytogenesis. All images show transverse spinal cord sections. **A-C**: Temporal expression profile of *sulf2* mRNA at E13.5 (A), E15.5 (B) and E17.5 (C). **D-D’**: Combined detection of *sulf2* (brown) and *fgfr3* (purple) mRNAs. D’ shows higher magnification of the area framed in D. **E**: Combined detection of *sulf2* (green) and *sox10* (red) mRNAs (red) viewed at high magnifications of E13.5 hemi-ventral spinal cord. **F-Q**: Detection and quantification of *fgfr3* mRNA+ parenchymal cells in wild-type (F, I, L, O) and *sulf2-/-* (G, J, M, P) embryos at E13.5 (F-H), E14.5 (I-K), E15.5 (L-M) and E18.5 (O-Q). Data are presented as mean ± SEM (***, p < 0.001 and *, p < 0.05). Scale bars = 100 μm.

These data not only reinforce the view that Olig2+ astrocytes depend on Sulf2 for their development but also highlight the specific role of Sulf2 for proper generation of this AP subtype.

## Discussion

### Two distinct types of Olig2+ glial precursors are generated in the developing spinal cord

Our work reveals a previously undefined heterogeneity within the population of spinal Olig2+ glial precursors generated at early gliogenic stages. So far, although pMN cells are recognized to generate some astroglial cells (Masahira et al., 2006; Tsai et al., 2012), OPCs were the only pMN-derived glial precursors to have been characterized. Typical OPC molecular identity relies on expression of Olig2 together with Sox10, Olig1 and PDGFRα, a set of genes activated as soon as pMN cells commit to the OPC fate (Bergles and Richardson, 2015). We identified a subtype of Olig2+ cells that do not activate this OPC program but, instead, adopt an AP molecular feature. Noticeably, OPCs (Olig2+/Sox10+) and APs (Olig2+/Aldh1L1+) are already distinguishable within the pMN domain, indicating that they segregate early, before leaving the progenitor zone. We did not detect real distinct subdomains within the pMN domain where the two Olig2+ cell subtypes appeared instead intermingled. This therefore questions the model of spatial segregation for glial-cell subtype specification in the developing spinal cord. Several different types of glial restricted precursors that can generate astrocytes and oligodendrocytes (O-A precursors) have been identified in the past, including GRP and O-2A cells (Liu and Rao, 2004). However, their existence in vivo has been the subject of debate (Rowitch and Kriegstein, 2010). Our results showing that OPCs and APs develop in the same region of the neural tube are consistent with the O-A precursor model, opening the possibility that Olig2+ pMN cells might share common features with these bipotent glial precursors at gliogenic stages.

### Sulf2, a specific player in regulating generation of Olig2-expressing APs

We show that lack of Sulf2 leads to defective Olig2+ AP generation but has no effect on OPC and Olig2 negative APs generation. How Sulf2 controls production of this AP subtype remains to be determined. Sulf2 is recognized as a critical regulator of signalling cues important in development, including FGFs, Wnt and BMPs (El Masri et al., 2017). Through its ability to catalyze specific 6-O-desulfation on HS chains, Sulf2 modulates signalling activities in many different ways, regulating either ligand secretion, processing and access to receptors (Rosen and Lemjabbar-Alaoui, 2010). Therefore, an attractive hypothesis could be that Sulf2 favors reception of an astroglial inductive signal in a subset of Olig2 progenitors. Our data, indicating heterogeneity in the levels of *sulf2* expression among pMN cells, are consistent with this hypothesis. However, the existence of such a signal remains so far elusive.

### Olig2 expression is retained in mature astrocytes of the ventral spinal cord

Our data show that a subset of ventral spinal cord astrocytes maintains expression of Olig2 until they reach their mature stage. Previous studies already noted the presence of Olig2-expressing astrocytes in the grey matter of the adult mouse spinal cord (Barnabé-Heider et al., 2010; Guo et al., 2011). Even if some Olig2+ astrocytes were occasionally observed in the white matter, we also found that most Olig2+ astrocytes are located in the grey matter, supporting the view they are related to protoplasmic astrocytes. Therefore, our work, by uncovering production of Olig2+ APs at initial stages of gliogenesis, sheds light on the developmental origin of these Olig2+ astrocytes. The most striking aspect is that ventral spinal Olig2+ APs maintain expression of Olig2 as they acquire a mature astrocytic identity. It is therefore obvious that, at least for this particular astrocyte subtype, Olig2 does not behave as a repressor of astrocytogenesis. Nonetheless, this observation raises the question as to whether expression of Olig2 confers specific functional properties to these astrocytes. During the past decade, evidence has accumulated that astrocytes represent remarkably heterogeneous cells that differ in their morphology, developmental origin, gene expression profile, physiological properties, function, and response to injury and disease (Ben Haim and Rowitch, 2017; Adams and Gallo, 2018). However, how the diverse astrocyte subtypes are generated remains poorly understood. Our study, by revealing the time of birth and molecular identity of pMN-derived astrocytes as well as a key regulator of their production during development, therefore represents an important advance in clarifying astrocyte heterogeneity in the spinal cord.

## Experimental procedures

### Mouse strains

All procedures were performed in agreement with the European Community guiding principles on the care and use of animals (Scientific Procedures) Act, 1987 and approved by the national Animal Care and Ethics Committee (APAFIS#01331.02) following Directive 2010/63/EU. Sulf2 mutant and Sulf2 floxed mice were genotyped as previously reported (Ai et al., 2007; Tran et al., 2012). The parental *olig2-cre+/−*;*sulf2fl/+;R26R-tomato* mice were generated by crossing heterozygous knock-in *olig2-cre* line (Dessaud et al., 2007) with mice that carried one floxed Sulf2 allele and the R26R-tomato reporter (*sulf2fl/+;R26R-tomato*). *olig2-cre+/−*;*sulf2fl/+;R26R-tomato* mice were crossed and *olig2-cre+/−*;*sulf2fl/fl; R26R-tomato* and *olig2-cre+/−*;*sulf2fl/+;R26R-tomato* littermate embryos were selected by genotyping. GFAP-GFP and Aldh1L1-GFP transgenic mice were obtained and genotyped as previously described (Nolte et al., 2001; Gong et al., 2003; Heintz, 2004). All mice were maintained on a C57BL/6 background.

### Thymidine analogue labelling

For proliferation assays, the timed pups (P0) were injected intraperitoneally with 5-ethylyl-2-deoxyuridine (EdU, Invitrogen) at 50 μg/g body weight during 2 days from day postnatal 0 to day postnatal 2. The animals were then sacrificed 2 hours after the last injection on day postnatal 2. EdU incorporation was detected using the AlexaFluor-555 Click-iT Imaging Kit (Invitrogen).

### Tissue processing

Spinal cord at the brachial level were isolated from E12.5 to E18.5 mouse embryos and fixed in 4% paraformaldehyde (PFA) in PBS overnight at 4°C. Adult mice were perfused intracardially with 4% paraformaldehyde (PFA, Sigma) in phosphate-buffered saline (PBS). Spinal cords were dissected and post-fixed in 4% PFA overnight at 4°C. Tissues were then sectioned either at 60–80 μm using a vibratome (Microm) or at 15 μm to 25 μm using a cryostat (Leica CM1950) after cryoprotection in 20% sucrose (Sigma) in PBS and freezing in OCT media on the surface of dry ice.

### In situ RNA hybridization and immunofluorescent staining

Simple or double in situ hybridization and immunofluorescent stainings were performed on transverse sections as previously described (Touahri et al., 2012; Ventéo et al., 2012). Digoxigenin- or Fluorescein-labeled antisens RNA probes for *sulf2*, a*ldh1L1, fgfr3, sox10* and *tenascin-C* were synthesized using DIG- or Fluorescein-labelling kit (Roche) according to the manufacturer’s instructions. Antibodies used in this study were as follows: goat anti-AldoC (SCB), rabbit and mouse anti-GFAP (DAKO), rabbit anti-NFIA (Active Motif), rabbit and goat anti-Olig2 (Millipore and R&D Systems), goat anti-Olig1 (R&D Systems), goat anti-Sox10 (Santa Cruz Biotechnology), rat anti-PDGFRα (BD Pharmingen), rabbit anti-Zeb1 (Novus) and mouse anti-Nkx6.1 (Hybridoma Bank). Alexa Fluor-594- or Alexa Fluor-488 or Alexa Fluor-647-conjugated secondary antibodies (Thermo Fisher Scientific) were used.

### Imaging, cell counting and statistical analysis

Images were collected from tissue sections using Leica SP5 or SP8 confocal microscopes. Images were processed using Adobe Photoshop CS6 and cell counts were performed using Image J software. Cell counts were performed on tissue slices harvested from level of the anterior limb (brachial spinal cord). Cell counts performed at embryonic and post-natal stages were done on hemi-sections of ventral spinal cord. The limit of the ventral region was defined by drawing a straight line through the midline dividing the spinal cord into two equal regions. For adult spinal cord, cell counts were limited to the ventral horns. Five sections per animal were quantified from at least five animals of each genotype. Data are presented as mean ± SEM and statistically analyzed using Student’s t test on Prism software (GraphPad).

## Acknowledgements

We especially thank X. Ai, N. Rouach, H. Kettenmann and B. Novitch for their generous sharing of Sulf2 mutant, Aldh1L1-GFP, GFAP-GFP and Olig2-Cre mice, respectively. We acknowledge A. Pattyn for providing valuable reagents and for critical reading of the mnauscript. We thank the ABC facility and ANEXPLO for housing mice and the Toulouse Regional Imaging Platform (TRI) for technical assistance in confocal microscopy. We acknowledge the Developmental Studies Hybridoma Bank, developed under the auspice of NICHD and maintained by the University of Iowa, Department of Biological Sciences, Iowa City, IA, for supplying monoclonal antibodies. Work in C.S.’ lab was supported by grants from ANR, ARC, ARSEP, CNRS and University of Toulouse. D. O. was supported by grants from ARSEP and ARC.

## Authors’ contribution

Conceptualization: D.O., C.D., B.G., C.S.; Methodology: D.O., N.E., P.C., Investigation: D.O., N.E., P.C., C.S.; Formal analysis: D.O.; Writing-Original Draft: D.O., C.S.; Supervision: C.S. Funding acquisition: D.O., C.S.

